# Metabolic plasticity supports a flexible nutritional symbiosis in *Cardiocondyla* ants

**DOI:** 10.1101/2025.10.17.683034

**Authors:** Phoebe Cunningham, José M Martín-Durán, Jan Oettler, Eva Shultner, Rayko Halitschke, Tobias Engl, Martin Kaltenpoth, Lee Henry

**Author notes:** **Correspondence and materials** requests should be addressed to Lee M. Henry. **Author contact information** Phoebe Cunningham José M Martín-Durán Jan Oettler Eva Shultner Rayko Halitschke Tobias Engl Martin Kaltenpoth Lee Henry.

## Abstract

Nutritional symbioses have repeatedly evolved in insects, yet how hosts regulate these partnerships to balance benefits across development and environments remains unclear. Ants provide an exceptional system to address this question, as several lineages maintain ancient symbionts that can be naturally lost without harming the host. Using *Cardiocondyla obscurior*, which carries and occasionally loses the vertically transmitted symbiont *Cand. Westeberhardia cardiocondylae*, we reveal the regulatory dynamics and function of this labile association. Symbionts provide shikimate-derived nutrients that enhance colony resilience under protein limitation, while hosts actively regulate symbiont abundance during protein scarcity and cuticle formation to optimise resource allocation. In the symbiont’s absence, ants compensate by upregulating genes enabling tyrosine acquisition from external sources. This metabolic flexibility allows colonies to exploit symbiont-derived and environmental nutrients, sustaining both growth and survival across nutritional contexts. Our findings reveal dynamic host regulation as a mechanism sustaining the persistence and adaptability of ancient symbioses.

## Introduction

Acquiring sufficient nutrients is a central challenge for insects, particularly for social species where dietary limitations can impact both individual and colony-level fitness. Nutrient-provisioning symbioses have repeatedly evolved as a solution, enabling insects to exploit nutrient-imbalanced diets and colonise specialised niches such as obligate sap- or blood-feeding^1–5^. However, symbioses are not without cost. They are metabolically demanding, and hosts must balance the energetic investment of maintaining symbionts against their nutritional benefits^6^. Studies have shown that insects can regulate symbionts through various mechanisms, such adjusting symbiont density, biasing them to specific developmental castes or sexes, or recycling them when their function is complete^7–11^. Yet, in most systems, how hosts regulate symbiont growth and coordinate metabolism to optimise benefits remains poorly understood.

Ants offer unique and underexplored models to address this question. At least four ant lineages have independently evolved intimate associations with *Sodalis*-allied bacteria— strictly vertically transmitted, intracellular symbionts housed in specialised host cells called bacteriocytes^12–14^. Despite their ancient origin and localisation within host-derived organs, we find these symbioses are remarkably labile: in some host lineages, symbionts are entirely absent from queens and even whole colonies in nature^12,13^. The evolutionary persistence of such flexible associations remains a puzzle. One hypothesis is that ants dynamically regulate their symbionts to meet nutritional demands, although the mechanisms by which this may occur remains unknown.

Genomic evidence suggests that *Sodalis*-allied symbionts in ants have all retained a conserved capacity to synthesise tyrosine via the shikimate pathway^13^. Tyrosine is essential for cuticle formation, melanisation, and immune function, but can only be obtained from protein (directly or via phenylalanine), making it limiting in protein poor niches such as arboreal habitats^15–17^. High carbohydrate to protein diets can also severely constrain ant colony growth^18^. We therefore hypothesise that, in environments with inconsistent protein availability, tyrosine is a key limiting resource for ants, selecting for alternative supplies of this amino acid, including symbiont-derived sources.

Here, we use the metabolically streamlined endosymbiont *Westeberhardia* in *Cardiocondyla obscurior* to uncover how hosts regulate symbionts in response to nutritional demands, and to confirm that the functional role of the symbiont is to maintain colony productivity when protein is limited. Arboreal *C. obscurior* provides an ideal model, combining experimental tractability with natural variation in symbiont presence in queens and entire colonies^12,19,20^. Through dietary manipulation and multi-omics analysis of symbiotic and aposymbiotic colonies, we show that *Westeberhardia* enhances colony fitness under protein scarcity via tyrosine and folate provisioning, and that the host flexibly regulates its metabolism and symbiont density in response to its diet. Our findings highlight the molecular basis of host– symbiont coordination and offer new insights into how a nutritionally important, yet evolutionarily labile, symbiosis can persist over millions of years.

## Results

### Symbiont improves colony productivity under protein starvation

To test the functional role of *Westeberhardia*, we simulated natural protein deprivation by feeding symbiotic and aposymbiotic *C. obscurior* colonies diets with reduced protein levels based on their standardised diet. Protein deprivation disproportionately affected later developmental stages in aposymbiotic colonies. Under complete protein starvation, pupal production in aposymbiotic colonies declined markedly, approaching significance (p = 0.07; Fig. S1A). This effect intensified at eclosion, where adult emergence dropped by 94% in aposymbiotic colonies compared to only 57% in symbiotic colonies (p = 0.02; Fig. 1A), underscoring the critical role of symbiont-derived resources in supporting development under protein scarcity. We also found that under optimal dietary conditions, symbiotic colonies were able to produce larger workers (*t*(22.1) = –2.3, *p* = 0.03; Fig. 1B) and exhibited a 75% lower risk of colony mortality, (*p = 0.01*, 95% CI: 0.081 – 0.79, Fig. 1C). Notably, worker cuticle thickness did not differ between groups, regardless of diet (Fig. S2). These results suggest *Westeberhardia* can aid in sustaining brood production during periods of protein deficiency, while maintaining colony health, and fortify worker size, under optimal conditions.

**Figure 1.**
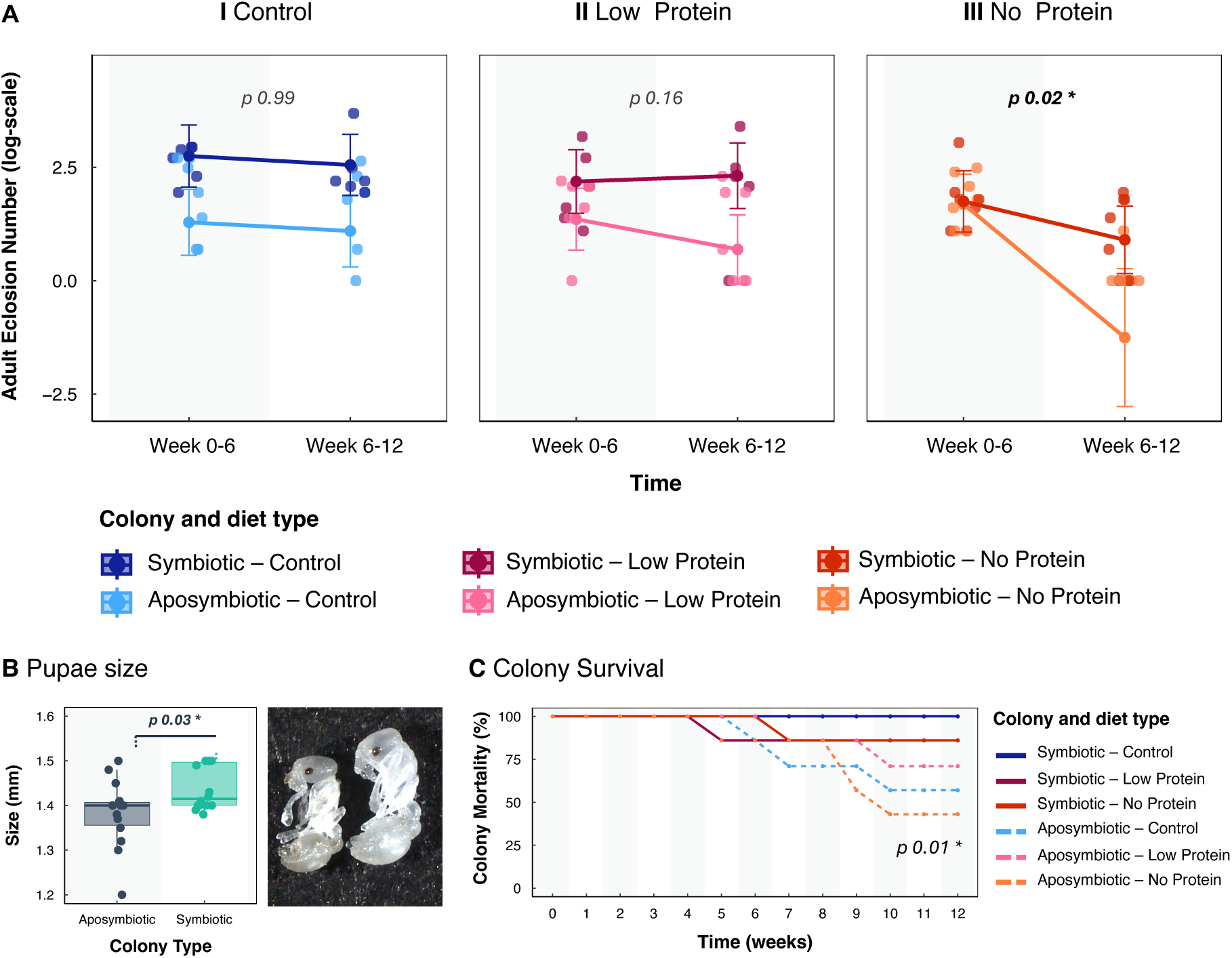
Effects of symbiont removal and protein starvation on colony and individual fitness. (A) Adult eclosion under control (dark/light blue), low-protein (dark/light pink), and no-protein (dark/light orange) diets in symbiotic vs. aposymbiotic colonies; (N = 7 per group, *p* values based on glmer modelling show significant differences between slopes); (B) Pupae size in symbiotic vs aposymbiotic colonies on control diets (N=13 per colony type); (C) Colony survival across protein treatments, *p* value based on Cox hazards testing represents significant difference between colony types. All model outputs are in Table S1.

### Symbiont density is dynamically regulated across host development and diet

We found that symbiont density varied dramatically across development, dropping sharply at the pre-pupal stage (∼day 19) after the ant has voided its meconium, which may disrupt gut-localised bacteriocytes (Fig. 2A). We then see a dramatic 1.6-fold recovery of symbiont density over 1-2 days when the pupa begins to form the morphology of an adult (Fig. 2A, days 19-28). This rapid rebound suggests a host-regulated mechanism that prioritises symbiont recovery during a key developmental transition when the adult cuticle is forming, a process the symbiont appears to be critical for.

**Figure 2.**
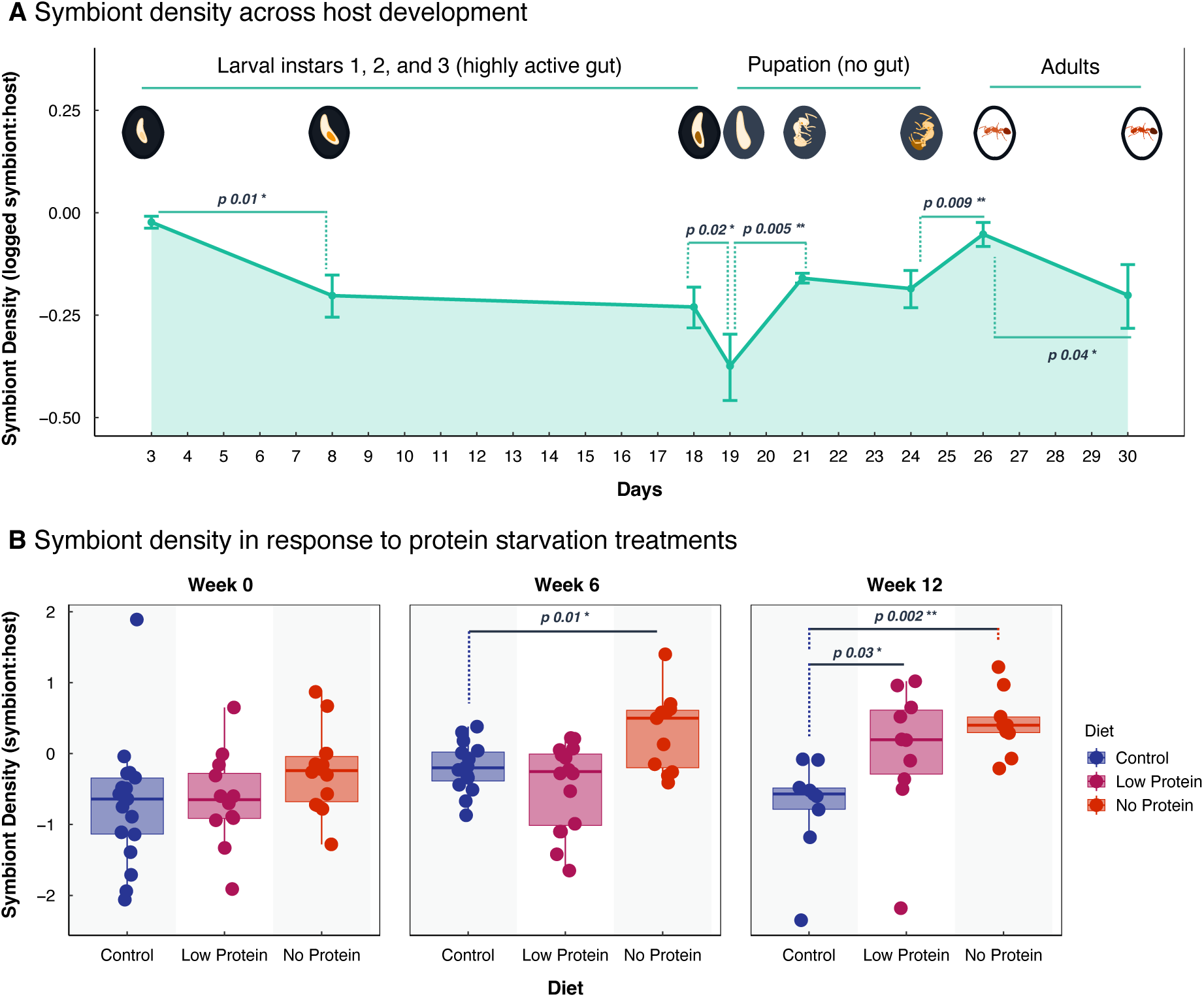
Symbiont density across host development and in response to protein starvation. Symbiont density, measured as the ratio of symbiont gene ribonucleoside-diphosphate reductase 1 subunit beta (nrdB) to host gene elongation factor-1 (EF-1) shown (A) across development stages (N=9 per stage); and (B) in response to no-protein (dark orange), low-protein (magenta), and control (blue) diets at weeks 0, 6 and 12 (N = 9–16).

We also found that symbiont density responded to dietary stress. Colonies deprived of protein showed significantly increased symbiont density by week 6 (*F* = 7.06, *df* = 39, no protein *p* = 0.02), and by week 12, colonies on both the low and no protein diets had significantly increased symbiont densities (*F* = 5.99, *df* = 25, low protein *p =* 0.03, no protein *p* = 0.002; Fig. 2B). These findings suggest an adaptive host response that increases symbiont abundance during key developmental stages and when exogenous protein is limited.

### Host transcription supports symbiont recovery at key developmental stages

Dual RNA-seq comparison of symbiotic versus aposymbiotic individuals (whole bodies) revealed over 650 differentially expressed genes (DEGs) at the pupal stage—far greater than at any other stage or caste (Fig. 3). By contrast, no DEGs were detected in *Westeberhardia* across host developmental stages (Fig. S3). These findings support our hypothesis that the symbiont plays a particularly important role during pupal development and that the principal method of symbiont regulation, e.g. population growth, is through host-derived mechanisms.

**Figure 3.**
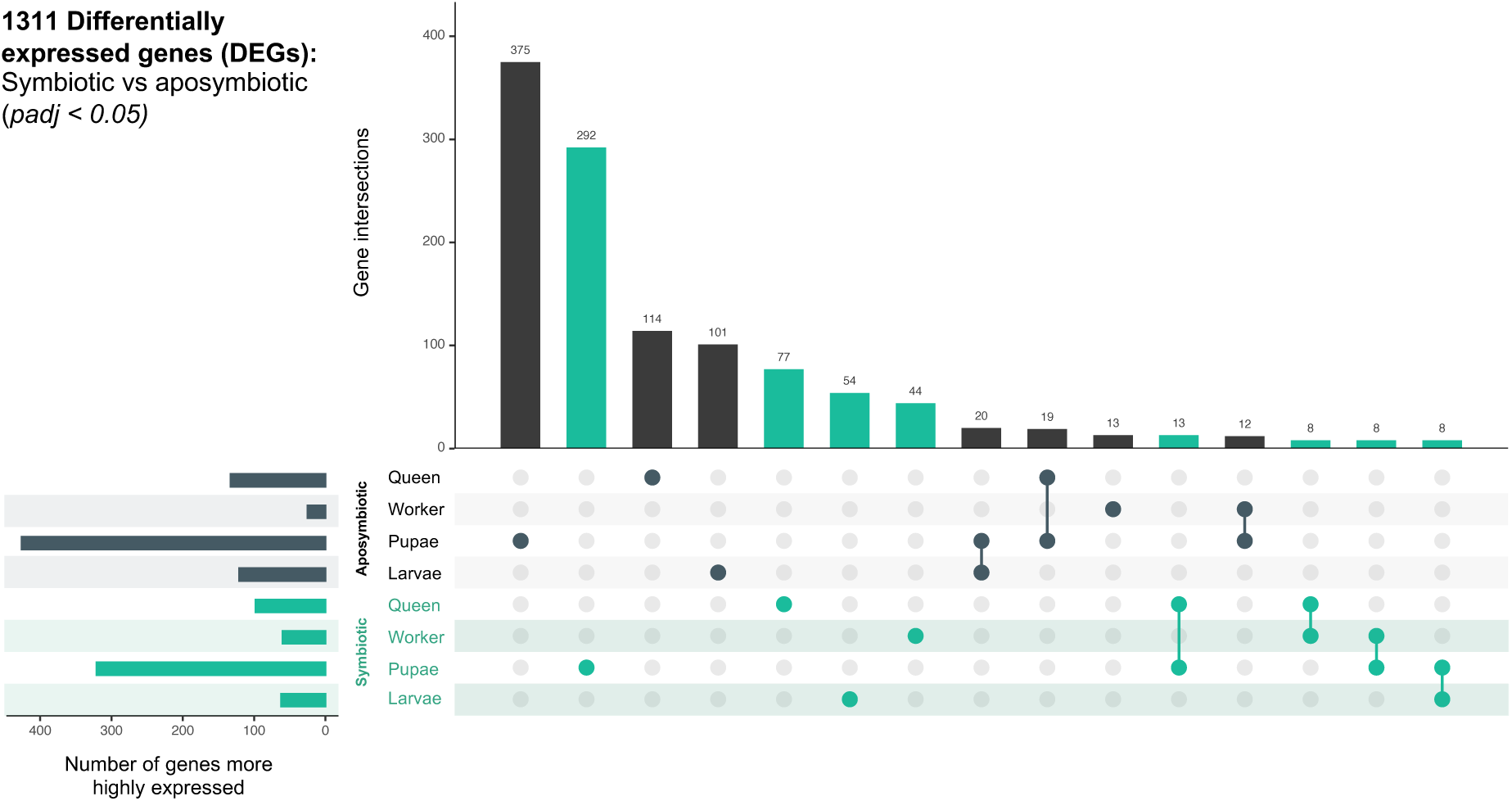
Differential gene expression between symbiotic and aposymbiotic colonies. Upset plot highlighting genes upregulated in symbiotic (turquoise) and aposymbiotic (grey) life stages or castes. Connected points indicate DEGs shared across multiple comparisons, representing a small subset of total DEGs.

Functional enrichment of DEGs revealed host genes likely co-opted to support symbiont recovery just before pupation, when symbiont function is most critical. Most notably, genes involved in sugar metabolism and transport were highly upregulated in symbiotic pupae compared to aposymbiotic individuals (Fig. 4A–B). This includes five genes that can break down sucrose, fructose, or lactose into β-D-galactose and β-D-glucose—substrates usable by *Westeberhardia—*coinciding with the rapid increase in symbiont titre (Fig. 4A). Several sugar transporters were also functionally enriched at this stage and more highly expressed in symbiotic pupae (Fig. 4B). These sugars could serve as an energy source by the symbiont but can also be fed directly into the shikimate pathway to produce 4-hydroxyphenylpyruvate, the molecular precursor that can be converted into tyrosine by the host (Fig. 4C). The sugar-metabolising gene myogenesis-regulating glycosidase was strongly expressed in symbiotic larval gut tissue (Fig. S4A–B). This suggests that hosts may increase carbohydrate intake during this life stage—similar to a mechanism in weevils that promotes endosymbiont proliferation^21^—then convert these into symbiont-compatible sugars before trafficking them to bacteriocytes.

**Figure 4.**
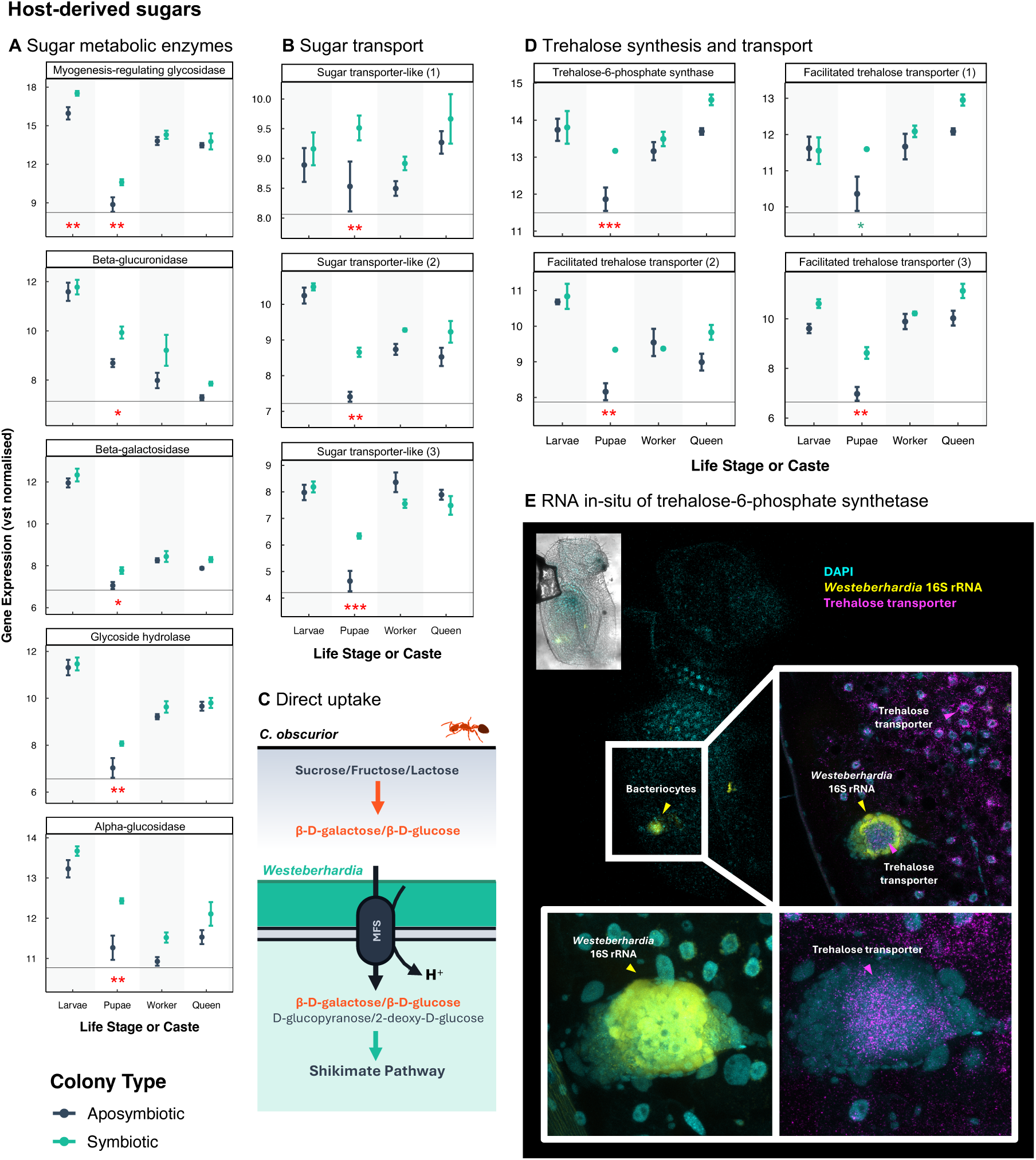
Expression patterns of putative host-symbiont regulatory genes. (A) Expression of host genes for enzymes to convert sucrose, fructose, or lactose into β-D-galactose/β-D-glucose; and (B) sugar transporters, are upregulated in symbiotic colonies; (C) Schematic showing *Westeberhardia* can uptake these sugars for use in the shikimate pathway; (D) Trehalose synthesis and transport genes show increased expression in symbiotic colonies; (E) Trehalose transporter shows concentrated expression in symbiotic larval bacteriocytes.

Gene expression was visualised using RNA-FISH with hybridization chain reaction (HCR): *Westeberhardia* was localised by tagging 16S ribosomal mRNA (yellow) using Alexa-514 fluorophore, and host trehalose-6-phosphate synthetase mRNA (magenta) using Alexa-647 fluorophore; host nuclei were stained with DAPI. Inset shows the same third-instar larva under brightfield with DAPI and RNA-FISH overlaid.

Our results also suggest symbiotic colonies may promote a more stable biochemical environment for *Westeberhardia* during pupation via trehalose-6-phosphate production and immune modulation. We find that trehalose-6-phosphate synthesis and transport genes were enriched in symbiotic pupae and localised to bacteriocytes (Fig. 4D–E). In the *Burkholderia*– bean bug symbiosis, trehalose is produced by the host to support symbiont colonisation by providing stress protection and metabolic stability^22,23^. Although we cannot rule out that *Westeberhardia* is using trehalose as an energy source, it lacks the trehalase genes needed to cleave it. Instead, trehalose may help stabilise the symbiont’s environment during periods of rapid physiological change. In addition, we observed reduced expression of genes regulating potassium ion channel activity in symbiotic colonies (Fig. S5), which play a role in immune defence in honey bees^24,25^. Additionally, genes involved in MAPK signalling and apoptosis were more highly expressed in the absence of the symbiont (Fig. S5), suggesting that *C. obscurior* may actively suppress its immune response when *Westeberhardia* is present. Similar immune modulation has been reported in other insect symbioses—cereal weevils use IMD pathway mediated genes to prevent endosymbiont escape from bacteriocytes and, contrastingly, aphids have entirely lost the IMD pathway, possibly in response to tolerance of *Buchnera aphidicola*^26,27^. Together, these findings suggest that during metamorphosis, a period of major physiological change, the host may downregulate immunity and promote a stabilising biochemical environment to facilitate symbiont recovery and re-establishment in developing bacteriocytes.

### Hosts adaptively regulate metabolism to meet tyrosine demands

Unlike many symbiotic insects with specialised diets, ants are omnivores and must flexibly manage nutrients from both symbiotic and exogenous sources. Remarkably, we find that *C. obscurior* meets this challenge by switching between two tyrosine biosynthesis pathways depending on whether the symbiont is present or not (Fig. 5A).

**Figure 5.**
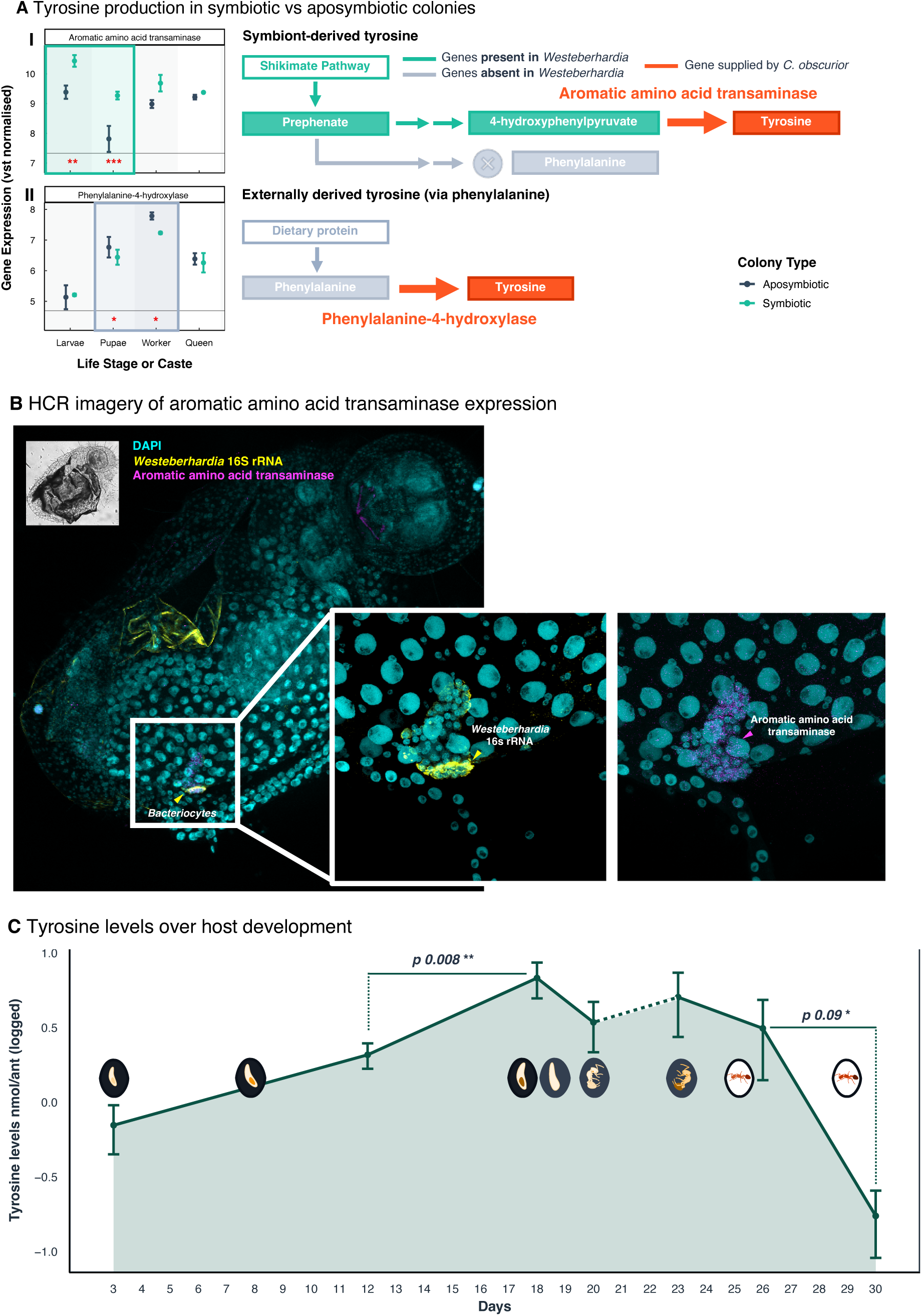
Flexible tyrosine biosynthesis strategies. (A) Expression of genes involved in two alternate pathways to produce tyrosine: (I) symbiont-derived 4-hydroxyphenylpyruvate converted by host aromatic amino acid transaminase (*tyrB*) (favoured in symbiotic larvae and pupae), and (II) host-derived conversion of dietary phenylalanine via phenylalanine-4-hydroxylase (*phhA*) (favoured in aposymbiotic pupae and workers); schematics indicate host vs. symbiont contribution; (B) *tyrB* expression localised to larval bacteriocytes using RNA-FISH via HCR (*Westeberhardia* 16S ribosomal mRNA in yellow; *C. obscurior tyrB* mRNA in magenta); (C) Tyrosine levels (nmol/ant) across development of symbiotic individuals; tyrosine is the only amino acid to increase during pupation (days 20–23, dotted line), when feeding has ceased (N = 9 per stage) (Fig. S6A).

When *Westeberhardia* is present, we find that host and symbiont metabolism align to biosynthesise tyrosine: the symbiont supplies 4-hydroxyphenylpyruvate via the shikimate pathway, while the host completes the process using its aromatic amino acid transaminase (*tyrB*). We find that *tyrB* is strongly expressed during larval and pupal stages in symbiotic colonies (Fig. 5A). Expression is also localised to bacteriocytes, indicating that the host is actively converting symbiont-derived 4-hydroxyphenylpyruvate on-site into tyrosine ahead of cuticle formation (Fig. 5B). Our metabolomic analysis also revealed that tyrosine was the only amino acid to increase during pupation - a stage when feeding ceases and no new protein is being ingested (Fig. 5C, S6A) - and to remain consistently elevated in symbiotic colonies under protein starvation (*F* = 6.6, *df* = 27, *p* < 0.02; Fig. S6B, Table S1). Additionally, expression of a cation transporter capable of shuttling L-DOPA, the tyrosine derivative needed for cuticle development, showed strong expression in bacteriocytes (Fig. S4C).

In contrast, when the symbiont is absent, we find the host relied on phenylalanine-4-hydroxylase (*phhA*) to convert dietary phenylalanine into tyrosine - an alternative pathway more highly expressed in workers and pupae lacking the symbiont (Fig. 5A, II). This metabolic switch highlights the host’s flexibility in maintaining tyrosine supply in the absence of the symbiont and suggests that symbiont loss may be tolerable in protein-rich environments, potentially explaining the natural variability in *Westeberhardia*’s presence across host populations. Together, these results show that coordinated host–symbiont gene expression promotes tyrosine and cuticle precursor synthesis under nutrient limitation, while host metabolic flexibility ensures these essential pathways are maintained in the symbiont’s absence.

### Cardiocondyla *exploits symbiont-derived folate provisioning when present*

We also found that genes involved in folate (vitamin B_9_) metabolism and transport were significantly enriched in symbiotic colonies, suggesting symbiont-derived folate is important for host development. Most notably, formyltetrahydrofolate synthetase is highly expressed in symbiotic larvae, pupae, and workers (Fig. 6A) and its expression is localised to bacteriocytes (Fig. 6B). This host enzyme is essential for carbon cycling and nucleotide synthesis, both critical for host growth and development. Folate provisioning by symbionts is common in blood-feeding insects^4,28–30^ and has also been documented in sap-feeding aphids^31^. *Westeberhardia* in *C. obscurior* has retained all genes for folate synthesis, however, some lineages of *Westeberhardia* have lost the capacity to produce folate, suggesting *Cardiocondyla* species may differ in their reliance on this as a symbiont-derived resource^13^.

**Figure 6.**
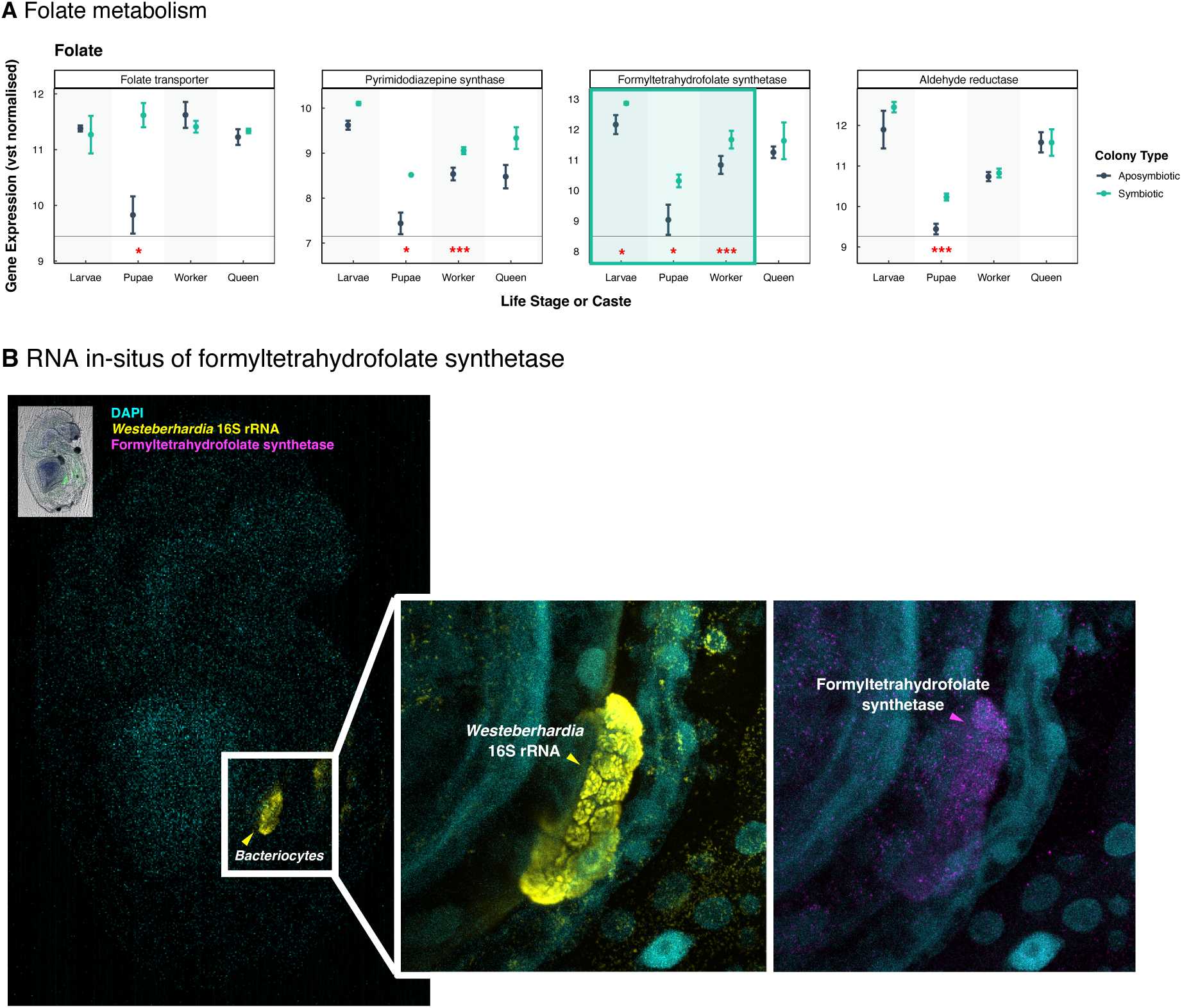
Folate provisioning in symbiotic colonies. (A) Host genes involved in folate metabolism, including formyltetrahydrofolate synthetase, a folate transporter and two genes involved in downstream processing, are more highly expressed in symbiotic colonies; (B) Formyltetrahydrofolate synthetase is shown to be localised to larval bacteriocytes using RNA-FISH via HCR hybridisation (*Westeberhardia* 16S ribosomal mRNA in yellow green; *C. obscurior* formyltetrahydrofolate synthetase mRNA in magenta).

## Discussion

### Environmentally responsive metabolism in insects

Metabolic plasticity is a key feature of insect adaptation, enabling responses to environmental cues, such as seasonality and diet, that have driven evolutionary innovations such as caste differentiation, diapause, and migration^32–36^. Our findings suggest that *Cardiocondyla obscurior* extends this plasticity to the regulation of its intracellular nutritional symbiont *Westeberhardia*. We show that ants can dynamically shift tyrosine biosynthesis between a symbiont-dependent route and a host-intrinsic pathway, allowing colonies to maintain fitness in the presence and absence of the symbiont.

Metabolic flexibility of this kind is likely rare in ancient intracellular symbioses, where hosts often become obligately reliant on symbiont-derived nutrients^6,37,38^. In contrast, *C. obscurior* has retained functional alternatives, activating symbiont-independent phenylalanine hydroxylation when *Westeberhardia* is absent. In most ancient nutritional symbioses, symbiont loss is accompanied by replacement that may result in some metabolic rewiring to accommodate the new symbiont^39^. True evolutionary losses of symbionts are relatively rare, but have occurred in hosts that transitioned to more nutrient-rich diets—scenarios that would also require the maintenance of pre-existing metabolic capacities to exploit these resources^9,40^. To our knowledge, this provides the first experimental evidence of a host mechanism that compensates for the loss of a strictly vertically transmitted nutritional symbiont, ensuring host survival. This metabolic versatility may reflect an evolutionary legacy of omnivory, which could favour the retention of redundant or alternative nutrient pathways. It raises the intriguing possibility that other ancient symbiotic omnivores, such as cockroaches^41^ and other ants^13^, may also preserve similar metabolic plasticity. However it may also be that the highly streamlined functional role of *Westeberhardia*, provisioning only shikimate-derived products, makes it more expendable in nature when compared to symbionts in omnivores with greater metabolic capacity—such as *Blochmannia* and *Blattabacterium*^41,42^.

### Host regulation and symbiont loss in nature

The facultative nature of *Westeberhardia* contrasts sharply with the widely held assumption that ancient, vertically transmitted nutritional endosymbionts are fixed and indispensable^13,37^. Our results support a model in which host control over symbiont density and function allows for conditional investment in the symbiosis, depending on nutritional context. Symbiont density increased under protein deprivation and during pupation. Transcriptomic data further support that this regulation is mediated almost entirely by the host, with no evidence of developmental stage-specific gene expression from the symbiont itself. Such host control may facilitate symbiont loss in environments where dietary protein is sufficient and may help explain why *Westeberhardia* is naturally absent in some *Cardiocondyla* queens and colonies^12,13^.

Similar losses occurs in other ant genera, such as *Formica*, where the symbionts presence varies within species despite long-term vertical transmission and bacteriocyte localisation^13^. Our findings lend empirical support to the hypothesis that the dietary flexibility associated with omnivory may reduce dependence on symbiont-derived nutrients and promotes the persistence of labile symbioses. Importantly, we find that *C. obscurior* can not only tolerate symbiont loss but can also actively reconfigure its metabolism in response—a key distinction from other systems where symbiosis breakdown can lead to fitness costs or mortality, with functional replacement often documented^43–46^.

### Nutritional symbiosis in a social context

In social insects, colony nutrient demands drive selection on endosymbiont maintenance and regulation in colony members^13,47^. We find that *Westeberhardia* contributes to colony-level fitness by increasing brood survival and adult emergence under protein stress, and by enhancing worker size even under optimal conditions. Worker numbers are positively correlated with queen production in *C. obscurior,* therefore investment in workers under protein stress may increase colony fecundity^48^. Notably, we observed no difference in cuticle investment across colony types, despite the link between symbiont-derived tyrosine and cuticle strength in other insects^45,49^. Tyrosine is required in high quantities during cuticle biosynthesis^16^. Our results suggest symbiont-derived tyrosine is essential for exoskeleton development and adult eclosion in *C. obscurior*. We hypothesise that tyrosine may be the key limiting nutrient in arboreal habitats - where *C. obscurior* nests and forages - rapidly compromising colony fitness when in short supply^15,50^. This could explain why symbiont-derived tyrosine appears to support reproduction, worker size, and colony growth in *C. obscurior* rather than investment in cuticle thickness.

Maintaining multi-queen colonies (polygyny), is often correlated with larger colony sizes and higher reproductive output^51^. Interestingly, all *Cardiocondyla* lineages that have undergone true evolutionary losses of *Westeberhardia* have are transitioned to single-queen (monogynous) colonies, whereas symbiont-retaining lineages have retained their ancestral polygynous state^13^. Given that *Westeberhardia* enhances brood production and survival, the symbiont may also play a role in maintaining this energetically demanding social strategy, particularly under protein-limited conditions. Moreover, several *Cardiocondyla* lineages, including *C. obscurior*, have spread globally through the tropics and sub-tropics and this invasive behaviour has only been observed in polygynous species^19^. The ability to modulate symbiont function in response to environmental conditions may buffer colonies against dietary variability, supporting traits commonly associated with invasion success, such as generalist diets and reproductive resilience^52–55^.

Our findings suggest that a context-dependent symbiosis contributes not only to nutritional homeostasis but may also enhance this species’ broader ecological plasticity. We propose that a flexible nutritional symbiosis - coupled with the ability to modulate it - may enhance this plasticity, allowing *C. obscurior* and possibly other *Cardiocondyla* species to thrive under variable or suboptimal conditions encountered during colonisation.

## Methods

### 1.0 Insect cultivation

#### 1.1 Colony collection and maintenance

*Cardiocondyla obscurior* colonies used in this study were originally collected in Japan in 2011 by Jan Oettler and colleagues^12^ and reared under controlled conditions at the University of Regensburg. In June 2022, six of these colonies were transferred to Queen Mary University of London, where they were maintained in a climate-controlled incubator replicating Regensburg conditions: a 12:12hr light:dark cycle with corresponding temperatures of 26°C (day) and 22°C (night), and 75% relative humidity. Colonies were fed three times per week with organic honey, cockroaches (once weekly), and fruit flies (twice weekly).

#### 1.2 Symbiont Removal and Validation

To remove *Westeberhardia*, *C. obscurior* colonies were heat-treated at 31°C for four weeks, maintaining all other conditions constant (adapted from Fan & Wernegreen, 2013)^56^. To confirm symbiont removal, DNA was extracted from larvae using the Qiagen DNeasy Blood & Tissue kit (QIAGEN Ltd., Manchester, UK), and screened via diagnostic PCR targeting the 16S rRNA gene (primers: WeBh_F2-for: 5′-CATTTGAATATGTAGAATGGACC-3′; WeBh-R2: 5′-AACTTTTACAAGATCGCTTCTC-3′). PCR conditions were: 95°C for 3 min; 39 cycles of 95°C for 5s, 60°C for 20s, 95°C for 10s; and a final elongation at 72°C for 5 min. Amplicons were run on 10% agarose gels; only colonies with no visible band were considered aposymbiotic. Larvae were re-sampled regularly to confirm the aposymbiotic state. Aposymbiotic colonies were rested for four weeks and allowed to re-queen before experimentation.

### 2.0 Gene expression analysis

#### 2.1 Sampling Design

Ants were flash-frozen and stored at –80°C. Samples from symbiotic and aposymbiotic colonies were pooled by life stage: (1) larval instars 1–3, (2) white and coloured pupae, (3) newly eclosed workers (identified by colour and mobility), and (4) queens. Samples within a treatment were pooled (N=10) to obtain sufficient RNA and homogenise potential variation among individuals.

#### 2.2 RNA Isolation, Library Prep, and Sequencing

Total RNA was extracted from whole bodies using TriReagent® and the Zymo Direct-zol® Mini Kit following manufacturers protocol. Samples were homogenised using pestles in 1.5ml RNase-free Eppendorf tube with liquid Nitrogen. RNA integrity was verified via Agilent TapeStation, and only samples with RIN scores >8.5 were used for sequencing. Due to low RNA yield in some samples, both TruSeq® Stranded Total RNA Library Prep Kit (Illumina) and the SMARTer® Stranded Total RNA-Seq Kit v2 (Takara) were used. Ribosomal RNA was depleted using both Illumina RiboZero Gold and NEBNext® rRNA Depletion Kits on all samples to maximise efficiency. Illumina sequencing, rRNA depletion and library preparation was performed by Macrogen (UK), generating ≥60 million paired-end reads per sample. Sample details can be found Table S2.

#### 2.3 Read processing

Reads were trimmed of the first 5 bases, adapters, and those with PHRED scores ≥25 using Trim Galore^57^, then checked with FastQC and MultiQC^58,59^. Residual rRNA was removed using SortMeRNA^60^. Reads were mapped to concatenated host and symbiont genomes using STAR aligner to maximise mapping fidelity^61–64^.

#### 2.4 Gene Expression Analysis

Raw STAR counts were analysed in R using DESeq2 (for differential gene expression) and limma (for batch correction via variance stabilising transformation)^65,66^. Pairwise comparisons between symbiotic and aposymbiotic life stages were conducted using lfcshrink, with significance defined as adjusted p < 0.05. Visualisations used ggplot2^67^. Genes of interest were identified through enrichment analyses of DEGs. To maximise detection of enriched genes, including poorly annotated but potentially important ones, we used multiple annotation tools—INTERPRO, Panther, GO, and KEGG—in combination with the universal enrichment function from the clusterProfiler^68–72^. Enriched genes and pathways were manually verified, and further annotated using NCBI’s BLAST^73^. Full gene expression data and annotations are in Tables S3–S5; code is in Supplementary Material.

#### 2.5 Gene expression localisation using Hybridisation Chain Reaction

RNA fluorescent in situ hybridisations (RNA-FISH) using hybridisation chain reaction (HCR) were used to validate patterns of spatial gene expression (adapted from Ferreira et al., 2021)^74^. Whole larvae were fixed in 37% paraformaldehyde/PBS/heptane (1:1:2) for 2-hours, then hybridised with gene-specific oPools Oligo Pools probes (from IDT) for 36–48 hours in a water bath at 37°C. Amplifier hairpins (Molecular Instruments) were applied overnight in the dark using a rocking table^75,76^, followed by staining of host tissue using DAPI diluted in PBS (10X) at a ratio of 1:1000 for 3 hours. Probes targeting *Westeberhardia* 16S rRNA (Alexa-514) were used to locate bacteriocytes; host genes were labeled with Alexa-647. Whole larvae were mounted on slides using glycerol and gene frames (Fisher Scientific, Leicestershire, UK, LE11 5RG). Imaging was conducted on a Leica Stellaris 8 confocal microscope and processed using Fiji (also known as ImageJ)^77^. Full HCR protocol is in the Supplementary Material.

### 3.0 Diet manipulations and phenotypic analysis

#### 3.0 Experimental Diets

To simulate dietary protein limitations, colonies were assigned to one of three diets: (1) control (insects and honey 3×/week), (2) low-protein diet (insects 1×/4 weeks, honey 3x/week), and (3) no-protein (honey only 3×/week). Prior to diet assays, each symbiotic colony was split in two; one half was heat-treated to remove the symbiont (Methods 1.2). After a four-week recovery, both symbiotic and now aposymbiotic colonies were each divided into three sub-colonies, followed by an additional two-week rest. Aposymbiotic status was regularly confirmed by PCR (Methods 1.2, Supplementary Material). Prior to treatment, all colonies were standardised with two mated queens and adjusted brood levels. Dead individuals were removed daily to reduce cannibalism risk. All colonies were handled identically.

#### 3.1 Fitness Assays

To assess pupae size, individuals were photographed blind and measured using a calibrated graticule. Pupal size differences were assessed with Welch’s t-tests^78^. Following dietary treatment initiation, brood production was recorded three times weekly using non-invasive stereomicroscopy. Every three weeks, samples were collected for: (1) cuticle analysis (newly eclosed workers), (2) metabolomics (late pupae), and (3) symbiont titre (third-instar larvae). Colony mortality was recorded throughout the experiment and analysed using Cox proportional hazards models with colony identity included as a random effect. For analysis of brood production we used generalised linear mixed effects modelling (glmer.nb from package lme4 in R), with a non-binomial distribution to compensate for overdispersion, and colony identity as a random effect^79^. All models tested for effects of symbiotic status (symbiotic vs. aposymbiotic), diet (control, low protein, no protein), and their interaction over time. Models were compared using likelihood ratio tests (LRT) and Akaike’s Information Criterion (AIC) and the R package emmeans was used to assess pairwise comparisons^80^. Visualisations were generated using *ggplot2*^67^. Fitness assay data is available in Table S6 and statistical model outputs in Table S1.

#### 3.2 Symbiont Density

Third-instar larvae (∼day 13) were flash-frozen and used for DNA extraction (Qiagen DNeasy kit). qPCR was performed using primers for *Westeberhardia* gene nrdB (ribonucleoside-diphosphate reductase 1 subunit beta)—5’ – GGA AGG AGT CCT AAT GTT GCG – 3’ and 5’ – AGC GTG CTC ACG AGT TTG TCC G – 3’—and host gene EF-1 (elongation factor 1-alpha 1)— 5’ –TCA CTG GTA CCT CGC AAG CCG A– 3’ and 5’ – ACC AGA AAT ATC TTT TGC ACG TT – 3’^64^. Each 10 µL reaction contained Lunar universal mix (NEB), 2 µL template, and 0.1 µL of each primer. Conditions: 95°C for 3 min; 39 cycles; melt curve 65–95°C in 0.5°C increments. qPCRs were run on 202 individuals in the same week with a single standard curve (values between 90-110% were accepted) (Table S7). Data were analysed with linear models (lme4) in R^79^, testing the difference in titre between colonies on different diets treatments at specific time points (week 0, week 6 and week 12). Results are in Table S1.

#### 3.3 Metabolomics

To measure amino acid levels, late-stage pupae were frozen, crushed under liquid nitrogen, and stored in 500 µL of 80% HPLC-grade methanol before being sent to the Max Planck Institute in Jena, Germany, for processing. Two negative controls were included. Due to sample limitations, amino acids were measured only in the control and low-protein diet treatments. Each 50 µL sample was diluted with 50 µL of an aqueous mix of ^13^C,^15^N-labelled amino acid standards (Aldrich, cat. no. 487910).

Samples were kept at 10°C, and 1 µL was injected into an UltiMate™ 3000RS UHPLC system with a Zorbax Eclipse XDB-C18 (3.0 × 50 mm, 1.8 µm) column, using acidified water and methanol in gradient mode^81^ (Schäfer et al., 2016). Natural amino acid isomers and internal standards were detected via EVOQ™ Elite triple-quadrupole MS (Bruker) with heated electrospray ionisation in positive mode (Spray voltage: 4500 V; Cone gas: 35 AU; Cone temp: 350°C; Probe gas: 50 AU; Probe temp: 500°C; Nebulizer gas: 60 AU). Samples were analysed in MRM mode, and concentrations were calculated using internal standards as per Schäfer et al. (2016)^81^. Post-run analysis was done using Bruker MS Workstation (v8.2.1). Generalised linear mixed effects models were used to test amino acid differences between colony types across diets, with colony identity as a random effect. Full calculations and concentrations are in Tables S8 and S9, model results are in Table S1.

#### 3.4 Cuticle thickness

Newly emerged workers were fixed in 4% formaldehyde in PBS, stored in ethanol, and sent to the Max Planck Institute (Jena, Germany). For micro-CT analysis, ants were fixed for 24 h in a 1:2 mix of 40% paraformaldehyde in PBS and 80% ethanol (Carl-Roth), washed in 100% ethanol, and then critical point dried (Leica EM CPD3000; 35 CO₂ cycles, low speed, 40°C). Samples were mounted and scanned using a Skyscan 1272 (Bruker; HAMAMATSU L10101-67 x-ray source, XIMEA xiRAY16 camera) at 1.000023 µm resolution, 0.105° rotation steps, 38 kV, 130 µA, 4× image averaging, 2×2 binning, 546 ms exposure.

3D datasets were reconstructed with NRecon (v2.1.0.1) using smoothing (1 iteration, kernel 2), ring artifact (0 or 5), and beam hardening correction (0–20%). Analysis in Dragonfly (v2022.1.0.1259, ORS) involved segmentation via upper OTSU threshold, followed by automated (close, fill X/Y/Z, erode, smooth; kernel ≤11) and manual refinement to isolate cuticle. Body and cuticle volumes were used to calculate proportional cuticular investment. Cuticle thickness meshes were generated with Laplacian smoothing (3 rounds) to extract average, median, and SD of thickness across the full body using Dragonfly. Full data are in Table S10.

## Supplementary Material

All supplementary material and sequencing data associated with this research will be available on publication.

## Author contributions

L.M.H conceived and designed the study. P.C reared the animals, collected the samples and performed all lab work carried out at QMUL, London. P.C performed all statistical analyses and generated all the visualisations. T.E performed the metabolomics work and R.H performed cuticle thickness measurements, both at the MPI, Jena. J.O and E.S provided the ants and expertise on their rearing and biology. J.M.M.-D consulted on the study design and provided expertise on RNA analysis and imaging work. M.K coordinated the technical lab support from MPI, Jena and provided expertise on the subject area. P.C and L.M.H drafted the manuscript and all authors read and commented on the manuscript.

## Acknowledgements

We thank all the members of the Henry and Martin-Duran lab for their support in the molecular work and statistical analysis, as well as the technical support received through M.K from the Max Planck Institute for Chemical Ecology, Jena. This work was performed using Queen Mary’s Apocrita HPC facility, which is supported by QMUL Research-IT (https://doi.org/10.5281/zenodo.438045). The research was funded by L.M.H.’s BBSRC-NSF/BIO (BB/W001632/1), Leverhulme Trust (RPG-2020-211) and BBSRC BB/W019698/1 grants.

## References

1. Douglas, A. E. Nutritional Interactions in Insect-Microbial Symbioses: Aphids and Their Symbiotic Bacteria *Buchnera*. Annu. Rev. Entomol. 43, 17–37 (1998).

2. Kirsch, R. et al. Symbiosis and horizontal gene transfer promote herbivory in the megadiverse leaf beetles. Current Biology 35, 640–654.e7 (2025).

3. McCutcheon, J. P., McDonald, B. R. & Moran, N. A. Convergent evolution of metabolic roles in bacterial co-symbionts of insects. Proc. Natl. Acad. Sci. U.S.A. 106, 15394– 15399 (2009).

4. Bing, X. et al. Unravelling the relationship between the tsetse fly and its obligate symbiont *Wigglesworthia* : transcriptomic and metabolomic landscapes reveal highly integrated physiological networks. Proc. R. Soc. B. 284, 20170360 (2017).

5. Moran, N. A., Tran, P. & Gerardo, N. M. Symbiosis and Insect Diversification: an Ancient Symbiont of Sap-Feeding Insects from the Bacterial Phylum *Bacteroidetes*. Appl Environ Microbiol 71, 8802–8810 (2005).

6. Bennett, G. M. & Moran, N. A. Heritable symbiosis: The advantages and perils of an evolutionary rabbit hole. Proc. Natl. Acad. Sci. U.S.A. 112, 10169–10176 (2015).

7. Ferrarini, M. G. et al. Coordination of host and endosymbiont gene expression governs endosymbiont growth and elimination in the cereal weevil Sitophilus spp. Microbiome 11, 274 (2023).

8. Vigneron, A. et al. Insects Recycle Endosymbionts when the Benefit Is Over. Current Biology 24, 2267–2273 (2014).

9. Buchner, P. Endosymbiosis of Animals with Plant Microorganisms. New York: Interscience (1965).

10. Benjamino, J. & Graf, J. Characterization of the Core and Caste-Specific Microbiota in the Termite, Reticulitermes flavipes. Front. Microbiol. 7, (2016).

11. Fukatsu, T. & Ishikawa, H. Soldier and male of an eusocial aphid, Colophina arma, lack endosymbiont: Implications for physiological and evolutionary interaction between host and symbiont. Journal of Insect Physiology 38, 1033–1042 (1992).

12. Klein, A. et al. A novel intracellular mutualistic bacterium in the invasive ant Cardiocondyla obscurior. ISME J 10, 376–388 (2016).

13. Jackson, R. et al. Convergent evolution of a labile nutritional symbiosis in ants. The ISME Journal 16, 2114–2122 (2022).

14. Sauer, C., Stackebrandt, E., Gadau, J., Hölldobler, B. & Gross, R. Systematic relationships and cospeciation of bacterial endosymbionts and their carpenter ant host species: proposal of the new taxon Candidatus Blochmannia gen. nov. International Journal of Systematic and Evolutionary Microbiology 50, 1877–1886 (2000).

15. Davidson, D. W. The role of resource imbalances in the evolutionary ecology of tropical arboreal ants. Biological Journal of the Linnean Society 61, 153–181 (1997).

16. Andersen, S. O. Insect cuticular sclerotization: A review. Insect Biochemistry and Molecular Biology 40, 166–178 (2010).

17. Moriyama, M. & Fukatsu, T. Host’s demand for essential amino acids is compensated by an extracellular bacterial symbiont in a hemipteran insect model. Front. Physiol. 13, 1028409 (2022).

18. Gutiérrez, Y. et al. Growth and survival of the superorganism: Ant colony macronutrient intake and investment. Ecology and Evolution 10, 7901–7915 (2020).

19. Heinze, J., Cremer, S., Eckl, N. & Schrempf, A. Stealthy invaders: the biology of Cardiocondyla tramp ants. Insect. Soc. 53, 1–7 (2006).

20. Seifert, B. The Ant Genus Cardiocondyla (Hymenoptera: Formicidae): The Species Groups with Oriental and Australasian Origin. Diversity 15, 25 (2022).

21. Dell’Aglio, E. et al. Weevil Carbohydrate Intake Triggers Endosymbiont Proliferation: A Trade-Off between Host Benefit and Endosymbiont Burden. mBio 14, e03333–22 (2023).

22. Elbein, A. D. New insights on trehalose: a multifunctional molecule. Glycobiology 13, 17R – 27 (2003).

23. Lee, J., Jeong, B., Bae, H. R., Jang, H. A. & Kim, J. K. Trehalose Biosynthesis Gene *otsA* Protects against Stress in the Initial Infection Stage of *Burkholderia* -Bean Bug Symbiosis. Microbiol Spectr 11, e03510–22 (2023).

24. O’Neal, S. T., Swale, D. R. & Anderson, T. D. ATP-sensitive inwardly rectifying potassium channel regulation of viral infections in honey bees. Sci Rep 7, (2017).

25. Fellows, C. J., Simone-Finstrom, M., Anderson, T. D. & Swale, D. R. Potassium ion channels as a molecular target to reduce virus infection and mortality of honey bee colonies. Virol J 20, (2023).

26. Nichols, H. L., Goldstein, E. B., Saleh Ziabari, O. & Parker, B. J. Intraspecific variation in immune gene expression and heritable symbiont density. PLoS Pathog 17, e1009552 (2021).

27. Maire, J., Vincent-Monégat, C., Masson, F., Zaidman-Rémy, A. & Heddi, A. An IMD-like pathway mediates both endosymbiont control and host immunity in the cereal weevil Sitophilus spp. Microbiome 6, 6 (2018).

28. Hosokawa, T., Koga, R., Kikuchi, Y., Meng, X.-Y. & Fukatsu, T. *Wolbachia* as a bacteriocyte-associated nutritional mutualist. Proc. Natl. Acad. Sci. U.S.A. 107, 769–774 (2010).

29. Duron, O. et al. Tick-Bacteria Mutualism Depends on B Vitamin Synthesis Pathways. Current Biology 28, 1896–1902.e5 (2018).

30. Tobias, N. J., Eberhard, F. E. & Guarneri, A. A. Enzymatic biosynthesis of B-complex vitamins is supplied by diverse microbiota in the Rhodnius prolixus anterior midgut following Trypanosoma cruzi infection. Computational and Structural Biotechnology Journal 18, 3395–3401 (2020).

31. Blow, F. et al. B-vitamin nutrition in the pea aphid-Buchnera symbiosis. Journal of Insect Physiology 126, 104092 (2020).

32. Hahn, D. A. & Denlinger, D. L. Energetics of Insect Diapause. Annu. Rev. Entomol. 56, 103–121 (2011).

33. Wilson, E. B. The Evolution of Caste Systems in Social Insects. Proceedings of the American Philosophical Society 123, 204–210 (1979).

34. Corona, M., Libbrecht, R. & Wheeler, D. E. Molecular mechanisms of phenotypic plasticity in social insects. Current Opinion in Insect Science 13, 55–60 (2016).

35. Ma, Z., Guo, W., Guo, X., Wang, X. & Kang, L. Modulation of behavioral phase changes of the migratory locust by the catecholamine metabolic pathway. Proc. Natl. Acad. Sci. U.S.A. 108, 3882–3887 (2011).

36. Kipyatkov, V. E. Seasonal life cycles and the forms of dormancy in ants (Hymenoptera: Formicoidea). Acta Soc. Zool. Bohem. 65, 211–238 (2001).

37. Moran, N. A., McCutcheon, J. P. & Nakabachi, A. Genomics and Evolution of Heritable Bacterial Symbionts. Annu. Rev. Genet. 42, 165–190 (2008).

38. Fisher, R. M., Henry, L. M., Cornwallis, C. K., Kiers, E. T. & West, S. A. The evolution of host-symbiont dependence. Nat Commun 8, 15973 (2017).

39. Mao, M. & Bennett, G. M. Symbiont replacements reset the co-evolutionary relationship between insects and their heritable bacteria. The ISME Journal 14, 1384–1395 (2020).

40. Hansen, A. K., Argondona, J. A., Miao, S., Percy, D. M. & Degnan, P. H. Rapid Loss of Nutritional Symbionts in an Endemic Hawaiian Herbivore Radiation Is Associated with Plant Galling Habit. Molecular Biology and Evolution 41, msae190 (2024).

41. Sabree, Z. L., Kambhampati, S. & Moran, N. A. Nitrogen recycling and nutritional provisioning by *Blattabacterium*, the cockroach endosymbiont. Proc. Natl. Acad. Sci. U.S.A. 106, 19521–19526 (2009).

42. Feldhaar, H. et al. Nutritional upgrading for omnivorous carpenter ants by the endosymbiont Blochmannia. BMC Biol 5, 48 (2007).

43. Matsuura, Y. et al. Recurrent symbiont recruitment from fungal parasites in cicadas. Proc. Natl. Acad. Sci. U.S.A. 115, (2018).

44. Perkovsky, E. & Wegierek, P. Aphid– *Buchnera* –Ant symbiosis; or why are aphids rare in the tropics and very rare further south? Earth and Environmental Science Transactions of the Royal Society of Edinburgh 107, 297–310 (2016).

45. Anbutsu, H. et al. Small genome symbiont underlies cuticle hardness in beetles. Proc. Natl. Acad. Sci. U.S.A. 114, (2017).

46. Martin Říhová, J., Gupta, S., Darby, A. C., Nováková, E. & Hypša, V. *Arsenophonus* symbiosis with louse flies: multiple origins, coevolutionary dynamics, and metabolic significance. mSystems 8, e00706–23 (2023).

47. Jackson, R. et al. Evidence of phylosymbiosis in Formica ants. Front. Microbiol. 14, 1044286 (2023).

48. Jaimes-Nino, L. M., Heinze, J. & Oettler, J. Late-life fitness gains and reproductive death in Cardiocondyla obscurior ants. eLife 11, e74695 (2022).

49. Kiefer, J. S. T. et al. Cuticle supplementation and nitrogen recycling by a dual bacterial symbiosis in a family of xylophagous beetles. The ISME Journal 17, 1029–1039 (2023).

50. Zuanon, L. A., Leão, R. E. O. S., Quero, A., Neves, K. C. & Vasconcelos, H. L. Nutrient Supplementation to Arboreal Ants: Effects on Trophic Position, Thermal Tolerance, Community Structure and the Interaction with the Host-Tree. Diversity 15, 786 (2023).

51. Boulay, R., Arnan, X., Cerdá, X. & Retana, J. The ecological benefits of larger colony size may promote polygyny in ants. J of Evolutionary Biology 27, 2856–2863 (2014).

52. Eyer, P.-A. & Vargo, E. L. Breeding structure and invasiveness in social insects. Current Opinion in Insect Science 46, 24–30 (2021).

53. Slatyer, R. A., Hirst, M. & Sexton, J. P. Niche breadth predicts geographical range size: a general ecological pattern. Ecology Letters 16, 1104–1114 (2013).

54. Snyder, W. E. & Evans, E. W. Ecological Effects of Invasive Arthropod Generalist Predators. Annu. Rev. Ecol. Evol. Syst. 37, 95–122 (2006).

55. Shik, J. Z. & Dussutour, A. Nutritional Dimensions of Invasive Success. Trends in Ecology & Evolution 35, 691–703 (2020).

56. Fan, Y. & Wernegreen, J. J. Can’t Take the Heat: High Temperature Depletes Bacterial Endosymbionts of Ants. Microb Ecol 66, 727–733 (2013).

57. Krueger, F. Trim Galore!: A wrapper around Cutadapt and FastQC to consistently apply adapter and quality trimming to FastQ files, with extra functionality for RRBS data. Babraham Institute (2015).

58. Ewels, P., Magnusson, M., Lundin, S. & Käller, M. MultiQC: Summarize analysis results for multiple tools and samples in a single report. Bioinformatics 32, 3047–3048 (2016).

59. Andrew, S. FastQC: A Quality Control Tool for High Throughput Sequence Data [Online]. (2010).

60. Kopylova, E., Noé, L. & Touzet, H. SortMeRNA: Fast and accurate filtering of ribosomal RNAs in metatranscriptomic data. Bioinformatics 28, 3211–3217 (2012).

61. Dobin, A. et al. Mapping RNA-seq with STAR. Curr Protoc Bioinformatics 51, 586–597 (2016).

62. Schrader, L. et al. Transposable element islands facilitate adaptation to novel environments in an invasive species. Nature Communications 5, 1–10 (2014).

63. Jackson, R. et al. Convergent evolution of a labile nutritional symbiosis in ants. The ISME Journal 1–9 (2022) doi:10.1038/s41396-022-01256-1.

64. Klein, A. et al. A novel intracellular mutualistic bacterium in the invasive ant Cardiocondyla obscurior. ISME Journal 10, 376–388 (2016).

65. Love, M. I., Huber, W. & Anders, S. Moderated estimation of fold change and dispersion for RNA-seq data with DESeq2. Genome Biology 15, 1–21 (2014).

66. Ritchie, M. et al. limma powers differential expression analyses for RNA-sequencing and microarray studies. Nucleic Acids Research 43, (2015).

67. Wickham, H. ggplot2: Elegant Graphics for Data Analysis. Springer-Verlag New York (2016).

68. Yu, G., Wang, L., Han, Y. & He, Q. clusterProfiler: an R package for comparing biological themes among gene clusters. OMICS: A Journal of Integrative Biology 5, 284– 287 (2012).

69. Paysan-Lafosse, T. et al. InterPro in 2022. Nucleic Acids Research 51, D418–D427 (2023).

70. Thomas, P. D. et al. PANTHER: Making genome-scale phylogenetics accessible to all. Protein Science 31, 8–22 (2022).

71. Aleksander, S. A. et al. The Gene Ontology knowledgebase in 2023. Genetics 224, 1–14 (2023).

72. Ashburner, M. et al. Gene Ontology: tool for the unification of biology. Nature Genetics 25, 25–29 (2000).

73. Camacho, C. et al. BLAST+: architecture and applications. BMC Bioinformatics 10, (2009).

74. A. G. Ferreira, A., Sieriebriennikov, B. & Whitbeck, H. HCR RNA-FISH protocol for the whole-mount brains ofDrosophila and other insects v1. Preprint at 10.17504/protocols.io.bzh5p386 (2021).

75. Dirks, R. M. & Pierce, N. A. Triggered amplification by hybridization chain reaction. Proc. Natl. Acad. Sci. U.S.A. 101, 15275–15278 (2004).

76. Schwarzkopf, M. et al. Hybridization chain reaction enables a unified approach to multiplexed, quantitative, high-resolution immunohistochemistry and *in situ* hybridization. Development 148, dev199847 (2021).

77. Schindelin, J., et al. Fiji: an open-source platform for biological-image analysis. Nat Methods 9, 676–682 (2012).

78. Zhenqiu Laura Lu & Ke-Hai Yuan. Welch’s t test. Preprint at 10.13140/RG.2.1.3057.9607 (2010).

79. Bates, D., Mächler, M., Bolker, B. & Walker, S. Fitting Linear Mixed-Effects Models Using lme4. Journal of Statistical Software (2015).

80. Searle, S. R., Speed, F. M. & Milliken, G. A. Population Marginal Means in the Linear Model: An Alternative to Least Squares Means. The American Statistician 4, 216–221 (1980).

81. Schäfer, M., Brütting, C., Baldwin, I. T. & Kallenbach, M. High-throughput quantification of more than 100 primary- and secondary-metabolites, and phytohormones by a single solid-phase extraction based sample preparation with analysis by UHPLC– HESI–MS/MS. Plant Methods 12, 30 (2016).

